# Pollinator traits and competitive context shape dynamic foraging behavior in bee communities

**DOI:** 10.1101/211326

**Authors:** Heather M. Briggs, Berry J. Brosi

## Abstract

Interspecific interactions (e.g. competition) can dynamically shape individual and species-level resource use within communities. Understanding how interspecific competition between pollinators species shapes resource use is of particular interest because pollinator foraging behavior (“floral fidelity”) is directly linked to plant reproductive function through the movement of conspecific pollen. Through targeted species removals, this study aims to gain a predictive understanding of how interspecific competition can influence pollinator foraging behavior. We explore how traits—specifically pollinator tongue length, known to dictate pollinator resource partitioning—influence behavioral plasticity and drive dynamic interspecific interactions. Our results demonstrate that bee species vary in their floral fidelity and that tongue length explains a large part of this variation. Bees with shorter tongues move between plant species (floral infidelity) more often than bees with longer tongues. We did not find significant variation in the response of bee species to a reduction in interspecific competition, but rather saw a guild-wide reduction in floral fidelity in response to the removal of the dominant bee species Finally, our results suggest that tongue length of the most abundant bee species, a site-level attribute, explains much of the site-to-site variation in pollinator foraging behavior. In particular, we found that as the tongue length of the most abundant bee in the site increases, the site level foraging fidelity decreases. With global pollinator populations on the decline, novel interactions between plants and pollinators are likely to occur. Exploring how the competitive landscape shapes foraging plasticity will help us generalize to other plant pollinator systems and begin to better predict the functional implications of competitive interactions.

## Introduction

Differences in traits among species may reduce interspecific competition and maintain diversity within a community (Grant 2006, Mayfield and Levine 2010). Generally, we still have a poor understanding of which traits influence the outcome of competition and community structure (McGill et al. 2006, Messier et al. 2010, HilleRisLambers et al. 2012). Traits such as body size (Wells 1988) or bill size/shape (Wiens and Rotenberry 1981, Grant 2006) and plant traits related to resource acquisition, such as root depth (Stubbs and Bastow 2004, Adler et al. 2010), can limit competition thus encouraging species coexistence. Our study explores the role of traits in mediating competition within a community context, via pollinator foraging behavior.

Pollinator foraging behavior directly affects plant reproductive output and ecosystem function through the transfer of pollen between plant individuals within a single foraging bout. If pollinators move between different plant species they can transfer heterospecific pollen, potentially reducing reproductive output (Ashman and Arceo-Gomez 2013, Briggs et al. 2016). Given the functional significance of pollinator resource use, it is important to understand the factors driving pollinator foraging behavior. The plants on which pollinators forage is a function of a number of factors including innate and learned preference, morphological traits, as well as direct and indirect competition with other pollinators in the community (Pimm et al. 1985, Cnaani et al. 2006, Stang et al. 2009, Brosi and Briggs 2013).

Decades of research has demonstrated that interspecific competition can influence pollinator foraging behavior. Largely in line with expectations derived from ecological theory, the range of resources that a species utilizes contracts as the strength of interspecific competition increases (Morse 1977, Hubbell and Johnson 1978, Inouye 1978, Pimm et al. 1985, Bolnick et al. 2010, Brosi and Briggs 2013, Fründ et al. 2013). In one example, Pimm et al. (1985) found that in the presence of one dominant competitor species, two other hummingbird species spent more time at a less rewarding feeder. In contrast, *without* interspecific competition from the dominant hummingbird, individuals of the other two species visited a feeder with high sucrose concentrations. Brosi and Briggs (2013) found that after a release from interspecific competition, bumble bees decreased their ‘floral fidelity’: pollinators moved more often between plant species within a single foraging bout. These changes in foraging behavior were associated with a significant decrease in reproductive output in a common alpine plant species. Fründ et al. (2013) provide another example in which pollinators’ flower preferences can be flexible and depend on community context (i.e., interspecific competition with other pollinator species present). In simplified experimental plant-pollinator communities, as competition between pollinator species increased, species often reduced their niche overlap by shifting to new plant species, which resulted in increased reproduction across the plant community. Thus, we know bees respond to competition and often do so strongly; but we don't know if bee species vary in their response to competition in complex assemblages of bee species or what traits are important in determining how they will respond.

Bumble bee (*Bombus*) communities provide an excellent system in which to empirically explore how trait differences drive foraging plasticity in response to interspecific competition. *Bombus* assemblages are often species-rich, and sympatric species typically have substantial overlap in their life history requirements (Goulson et al. 2008). Furthermore, traits that affect resource acquisition and foraging efficiency can influence how species partition resources within a community (Abrams and Chen 2002, Grant 2006). Tongue length is a trait that directly determines which resources a bumble bee can access and how resource selection varies among species (Heinrich 1976, Inouye 1978, McGill et al. 2006, Stang et al. 2009). In general, long-tongued bumble bee foragers visit flowers with deep corollas and short-tongued bumble bees forage on shallow flowers (Heinrich 1976, Stang et al. 2006, McGill et al. 2006). Still, bumble bees are known to be labile in their foraging patterns if more rewarding floral resources become available or the competitive landscape shifts (Inouye 1978, Gegear and Thomson 2004, Brosi and Briggs 2013). Tongue length appears to be important for structuring foraging preferences but we are lacking experimental work that evaluates how traits such as tongue length influence pollinators’ response to competition.

We systematically manipulated interspecific competition in bumble bee communities through targeted single species removals and examined patterns of species-specific plasticity in resource use in the remaining pollinators. Specifically we examined to what extent tongue length explains species-specific difference in the pollinators’ foraging behavior in response to release from competition. We worked in spatially replicated plots in natural communitites varying in plant and bee community composition. Utilizing removals in these natural communitites allowed us to examine if the identity of the most abundant bee species (i.e. compeititve context) influences bee foraging behavior. We focused on the foraging response of the remaining bees in the community with respect to floral fidelity, or, within plant species movements within a single foraging bout. Floral fidelity is critical for many plants species’ reproductive success because transfer of conspecific pollen must occur in order for fertilization to take place.

We asked specifically (1) How do bee species vary in their overall patterns of floral fidelity? (2) To what extent does tongue length explain variation in species-specific foraging behavior? (3) How does the competitive context in which bees forage influence their floral fidelity, and is there a systematic relationship between the traits (i.e. tongue length) of the dominant bee species and the foraging patterns of the remainder of the bees at a given site?

## Methods

### Study Sites

We worked in 28 subalpine meadow sites in the landscape surrounding the Rocky Mountain Biological Laboratory (38° 57.5′N, 106°59.3′W, 2,900 m above sea level), in the Gunnison National Forest, western Colorado, United States. Each site consisted of a 20 × 20-m plot, all with the same dominant plant species (*Delphinium barbeyi*). A minimum distance of 1 km separated any two sites. We collected data over three summer growing seasons (June-August), in 2010, 2011 and 2013.

### Manipulations

We assessed each plot in a control state, waited one day, and then assessed each plot in a manipulated state. We kept the interval between control and manipulated states short because of the rapid turnover in flowering phenology in our high-altitude system, allowing us to keep the plant community constant in our control–manipulation comparisons (Langenheim 1962, Brosi and Briggs 2013). Manipulations reduce interspecific competition through the temporary, non-destructive removal of the most abundant bumblebee species in each plot. We determined the most abundant bee via inventory of *Bombus* species richness and abundance on the control day using nondestructive aerial netting, with two field team members netting for a 20-min period, not including handling time (the time from when a bee was in the net until it was in a closed vial). To avoid double-counting, we kept each bee in an individual glass vial, identified to species, and kept in a cool, dark cooler until the inventory time period was over, at which point bees were released.

On the manipulation day we removed the most abundant bee species (as determined two days previously in the control state). The removals were accomplished through targeted hand-netting, and we minimized disturbance of other bees and vegetation by carefully placing the insect net over entire inflorescence and allowing bee individuals to fly up into the net (Inouye 1978). Captured bees were transferred to vials and placed in a cooler during the manipulation and released unharmed afterward. We used as much time as necessary to remove essentially all individuals of the target species from the sample plot and immediately adjacent area (typically in 1-2 hours we would achieve ~98% removal). We left a period of at least 30 minutes between manipulative bee removals and subsequent sampling to minimize the impact of the disturbance on the foraging activities of other bees. We recorded both the abundance of removed (captured) individuals, as well as the number of un-captured “escapees” of the most abundant species that were observed during bee sampling. We assessed resource use (plant species visited within a single foraging bout) in each site in both a control and a manipulated state. Each site was only used once in a manipulated and controlled state per year (i.e., no sites were re-sampled within a single season).

### Foraging observations

We directly followed the foraging sequences of *Bombus* individuals in both the control and manipulated states. We recorded the identity of each plant species visited in a foraging sequence. We discontinued an observation when the bee was lost from sight, when it ventured more than 5 m outside of the plot, when it had been observed for 10 full minutes, or when we had tallied 100 individual plants visited. We discarded observations of fewer than five plants visited. The number of individuals observed per/state/site varied due to bee abundance (mean = 33 individuals per site; range = 8–54).

### Tongue Length measurements

The data on proboscis lengths of workers was taken from published measurements of bumblebees collected on the Front Range of the Colorado Rocky Mountains and overlap with the sites and species that we used for this study (Inouye 1980). Measurements indicate the sum of the individual lengths of the prementum and glossa. The mean tongue length of each bumble bee species was assigned to each individual bee and used in the trait analysis ([see Table 1 for tongue lengths] Inouye 1978, 1980, Pyke 1982).

**Table 1:** Tongue lengths of sympatric *Bombus* spp. Mean and standard deviation of 50 individuals per species from Inouye (1976).

| *Bombus* Species | mean<br>mm | sd | N sites in<br>which sp. was<br>removed |
| --- | --- | --- | --- |
| *B. occidentalis* | 5.71 | 0.25 | 0 |
| *B. bifarius* | 5.75 | 0.37 | 6 |
| *B. sylvicola* | 5.79 | 0.58 | 1 |
| *B. flavifrons* | 7.31 | 0.8 | 9 |
| *B. balteatus* | 9.36 | 0.62 | 4 |
| *B. californicus* | 10.01 | 0.75 | 0 |
| *B. nevadensis* | 10.13 | 0.68 | 2 |
| *B. appositus* | 10.48 | 0.95 | 5 |

### Data analysis

#### Quantifying Floral fidelity

Floral fidelity was measured as the binomial counts of individual bee foraging movements that were conspecific (between individuals of the same plant species) vs. heterospecific (between individuals of different plant species) (Brosi and Briggs 2013). We used GLMMs with binomial errors using the logit link in the lme4 package for R (Hothorn et al. 2013) to model the floral fidelity response variable. Data from individual bees foraging within a site cannot be considered independent becase bees within a site are likely to be closely related genetically, and environmental conditions are similar; therefore, site was included as a random intercept term (Bolker et al. 2009). Relative to a binomial distribution, our data were overdispersed, which we corrected by including an individual-level (i.e. bee individual) random intercept term (Elston et al. 2001). We used the R statistical programming language (R Core Development Team 2012) for all models.

#### Species Specific differences

We first ran a null model (M0) that included only the two structural random effects: one to control for pseudoreplication for site and another to correct for over-dispersion in the binomial response variable (see above).

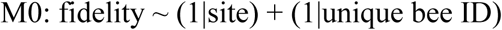

These structural terms were retained in all models. We then added state (control or manipulation) as a fixed effect to M0, giving M1 below, to assess whether the removal of the most abundant bee had a guild-wide impact on foraging fidelity.

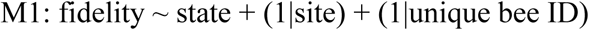

We compared M1 and M0 through a likelihood ratio test (LRT) and retained state as a fixed effect in all subsequent models (see Results, Table 2). Next, we estimated the magnitude of between-bee species variation in foraging dynamics by adding bee species as a random intercept (model M2) or as a random intercept and a random slope (model M2b) with respect to state to model M1:

**Table 2:** Results of generalized linear mixed-effects models with binomial errors *P < 0.05; **P < 0.01; ***P < 0.001; TL = tongue length MTL = manipulated tongue length (i.e. tongue length of the species removed from each site); R^2^C = describes the proportion of variance explained by both the fixed and random factors; R^2^M describes the proportion of variance explained by the fixed factors alone; LRT = likelihood ratio test.

| model | Fixed effects |  |  | Random effects |  |  | Model Comparison |  |  |  |
| --- | --- | --- | --- | --- | --- | --- | --- | --- | --- | --- |
|  | state | TL | MTL | Species<br>α/β | Site | Individual | AIC | R<sup>2</sup>C | R<sup>2</sup>M | LRT |
| MO | -- | -- | -- | -- | 0.880 | 5.240 | 2079 | 0.093 | 0.000 | -- |
| M1 | 0.93*** | -- | -- | -- | 0.930 | 5.000 | 2064 | 0.122 | 0.022 | 0:1, $p = 3.2 \times 10^{-5}$ |
| M2 | 0.72** | -- | -- | 0.444 | 1.159 | 3.900 | 2036 | 0.199 | 0.021 | 1:2, $p = 4.5 \times 10^{-8}$ |
| M2b | 0.726** | -- | -- | 0.10/0.24 | 1.115 | 3.959 | 2038 | 0.199 | 0.022 | 2:2b, $p = 0.319$ |
| M3 | 0.899*** | -0.274** | -- | 0.160 | 0.920 | 4.170 | 2034 | 0.176 | 0.055 | 2:3, $p = 0.035$ |
| M4 | 0.937*** | -0.389*** | 0.301*** | 0.000 | 0.178 | 6.404 | 2025 | 0.127 | 0.112 | 3:4, $p = 0.003$ |

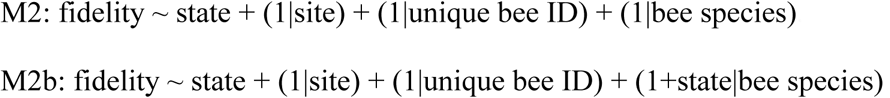

These models allowed us to estimate how much bee species differ in both their base-line fidelity (i.e. random intercept) and in their response to the manipulation (i.e. random slope) by estimating a variance term for each. To test if the random effects associated with bee species improved the model fit, we compared models M1 and M2, as well as M2b and M2, using LRT and by computing differences in the values of Akaike's Information Criterion (∆AIC) between the models.

#### Traits

To examine if bee species-level traits explain the species-specific differences in foraging behavior, we added tongue length as a continuous fixed factor to M2 yielding M3 below.

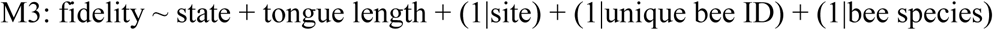

We then asked whether tongue length explains differences in base-line floral fidelity among bee species. We did so by first comparing M3 to M2 via LRT and by calculating ∆AIC between these models to determine the overall impact on the model of adding tongue length as a fixed effect. If tongue length explained base-line differences in foraging behavior that was otherwise modeled as a random effect, the variance estimate for bee species intercepts should be reduced in M3 relative to M2. We calculate this relative reduction in variance as

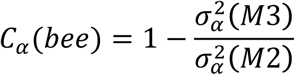

following Jamil (2013), where subscript α indicates random intercept terms. The numerator indicates the “residual” variation among bee species after taking account of tongue length (M3), while the denominator is the ‘total’ variance among bee species from the model without tongue length (M2). *C*_*α*_ > 0 suggests inter-specific differences in base-line fidelity may be explained by differences in their tongue lengths.

We evaluated variability in the estimates for 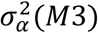 and 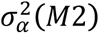—and thus, the difference between them—through parametric bootstrapping, i.e. by simulating data from the maximum likelihood fits of M3 and M2 and then re-fitting the model. This approach generates a set of coefficient fits for 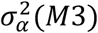 and 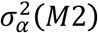. We then compared the bootstrap distributions for the varience terms using the non-parametric two-sample Kolmogorov–Smirnov (KS) test, which examines the null hypothesis that the boot strap replicates for the variance terms are not different at the *p*≤0.05 level.

#### Site level attributes

Finally, we explored if the community in which bees are locally embedded influenced their foraging patterns. Specifically, we asked whether site-level variance in base-line foraging patterns of the observed bees is a function of the identity of the most abundant bee species at that site. We consider this a site-level attribute and a stand-in for competitive context for the observed bees. To do so, we added a term to M3 for the tongue length of the bee species that was the most locally abundant at the site, and thus the bee that we non-destructively removed from the site in our manipulations (‘manip. tongue length’), as a continuous fixed factor, giving M4 below:

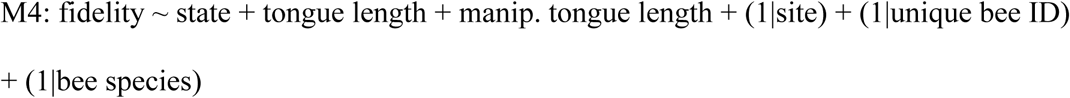

To assess overall effects on model fit of adding this additional term, we compared M4 to M3 using LRT and by calculating ∆AIC. We compared variance estimates for the site intercept from M4 to that from M3 by calculating *C*_*α*_ (*site*), with site variance from M4 in the numerator and that from M3 in the denominator. This comparison examines to what extent the trait of the most abundant species at a site captures site-to-site variance in baseline fidelity that would otherwise be contained within the random intercept term for site. Again, a result of *C*_*α*_ > 0 indicates that the traits of locally abundant bee species explain site-to-site variance (Jamil 2013). The distributions for 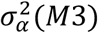 and 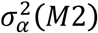 for site were compared using a KS test.

## Results

### Overall effect of manipulation

The addition of manipulated state as a fixed effect in M1 confirmed that the removal of the most abundant bee resulted in a guild-wide reduction in foraging fidelity when examined across the three year study period (Coeff = 0.93, SE = 0.22, *p* = 3.87 × 10^−5^; Table 2; Fig. 1), consistent with previous work that analyzed only the first two years of this dataset (Brosi and Briggs 2013). Furthermore, we found that M1 was a significantly better model when compared to the M0 (∆AIC = −15.29, LRT: *p* = 3.2 × 10^−5^), we therefore retained state as well as the structural random effects (site and bee individual) in all subsequent models.

**Figure 1:**
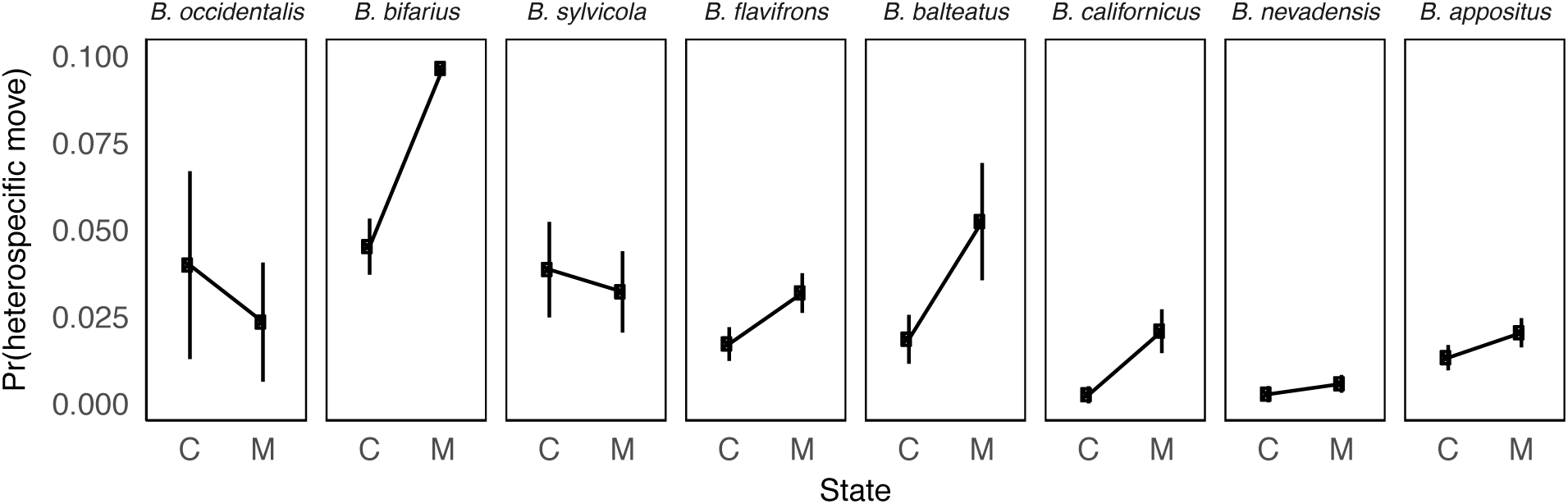
Species specific response to pollinator removals. **A.** Mean effect of experimental state (‘C’ = control, ‘M’ = manipulation) on the probability that foraging moves are between heterospecific plant species, plotted for each bee species. Bee species panels are arranged left to right in increasing order of their tongue length. 95% CI around the mean heterospecific move probability calculated on the basis of pooled counts for each state for each bee species.

### Species-specific differences

Adding bee species as a random intercept (M2) and as a random slope term (M2b) to M1 significantly improved model fit over M1 (M2 vs. M1: ∆AIC = −28, LRT: *p* = 4.8 × 10^−7^; M2b vs. M1: ∆AIC = −26, LRT: *p* = 4.8 × 10^−7^). Model M2b revealed only slight interspecific variation in the *response* to the manipulation that was not already captured by the main effect of state (variation in the random slope term for bee species = 0.10, see Table 2). While M2b was a better model than M1, it did not improve model fit compared to M2 (∆AIC = 1.72, LRT: p > 0.5). In contrast, model M2 reveals substantial interspecific variation in the *baseline* floral fidelity of bees (variation in the random intercept term for bee species = 0.24; Table 2). Subsequent model comparisons were made with model M2. The fixed effect of state remained highly significant (Coeff = 0.72, SE = 0.26, *p* = 0.005, see Table 2) in this model.

### Traits

#### Baseline differences in species specific fidelity

The addition of the trait level fixed effect in M3 largely explained the *baseline* variation in inter-specific floral fidelity (when compared to M2). The baseline fidelity of bees with shorter tongues is lower than that of bees with longer tongues (i.e. short tongue bees move between plant species more often) (Fig. 2, S2). The variance estimate for the random intercept term for bee species dropped from 0.444 in M2 to 0.164 in M3, giving *C*_*α*_(*bee*). Furthermore, M3 is a significantly better model than M2 (∆AIC = 2, LRT: *p* = 0.043, Table 2), indicating the importance of tongue length as an explanatory factor for floral fidelity. The bootstrapped distribution of estimates for the bee species random effect from M3 substantially overlapped 0, while that for M2 was well above zero (Fig. S1). Further supporting our interpretation of this term, a KS-test of the difference between these bootstrapped distributions was highly significant (*p* < 10^−10^).

**Figure 2:**
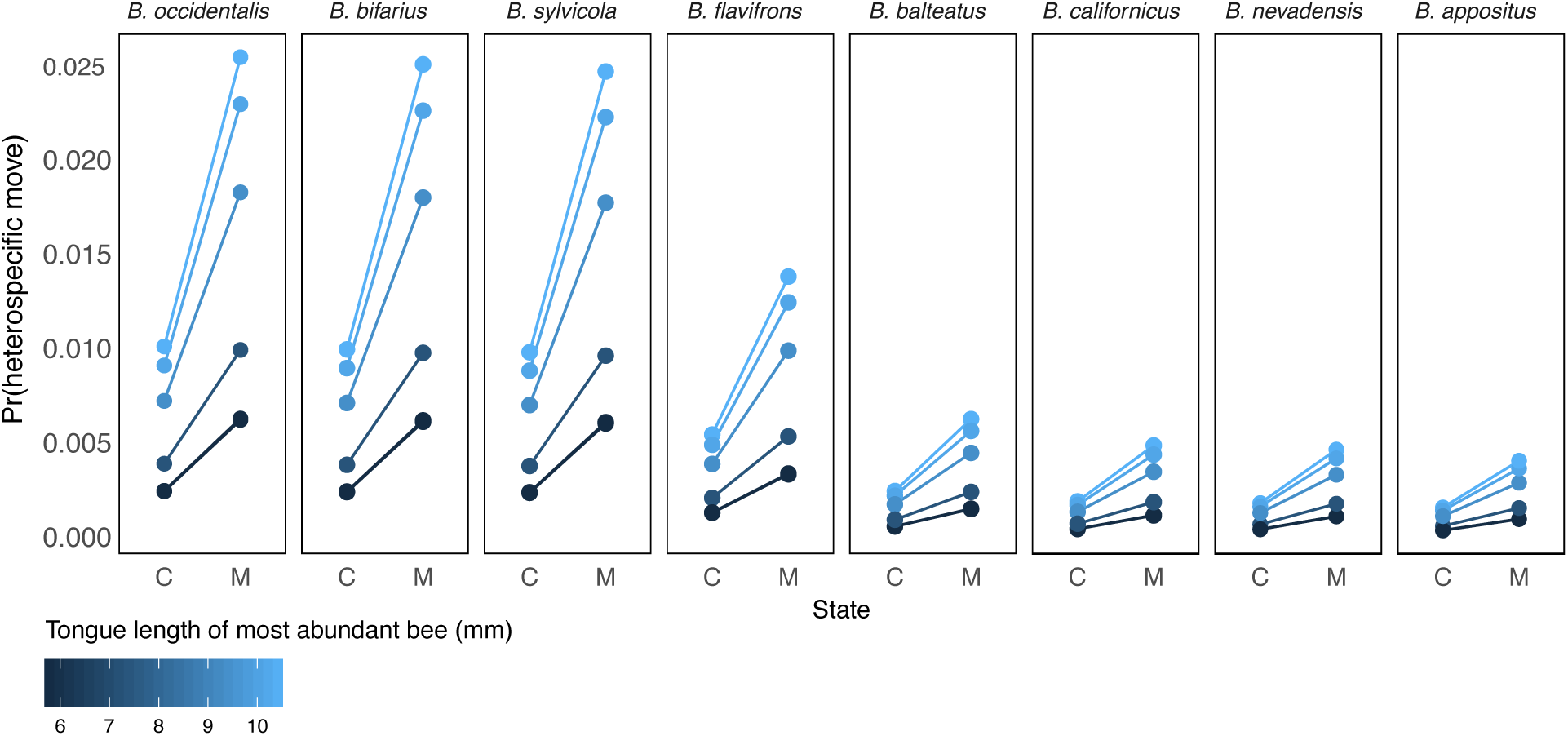
Predicted impacts of the fixed effects from model M4 on the probability of heterospecific foraging moves. Contribution of random effects to the variance in heterospecific moves are not displayed in order to reveal the how the three fixed effects (see Table 2) shape foraging fieldity. Shown are the predicted mean foraging behavior of each bee species in each state (‘C’ = control, ‘M’ = manipulation) across a gradient of sites where the manipulated bee varies in tongue length (‘tongue length of most abundant bee’). Bee species panels are arranged left to right as shortest to longest tongue length.

#### Site level attributes

We found that adding the tongue length of the removed bee as a fixed effect, (a site-level attribute), largely accounted for site-to-site variation in bees’ floral fidelity (over and above species-level differences) (∆AIC = –8.7; LRT: p = 0.001). That is, site level variation in bumble bee floral fidelity is largely explained by the tongue length of the most abundant bee species (i.e., the species removed experimentally): variance estimates for the site intercept dropped from 0.918 in M3 to 0.178 in M4, giving *C*_*α*_ = 0.80, suggesting that there is little variance left to explain after accounting for site level variation (Table 2). The three fixed effects, state, tongue length and the tongue length of the removed bee species, are also statistically significant in M4 (Coeff = 0.937, SE = 0.242, *p* = 0.0001, Coeff = 0.734, SE = 0.276, *p* = 7.46 × 10^−8^, Coeff = 0.301, SE = 0.081, *p* = 0.0001, respectively; Table 2, Fig. 2). Furthermore, the bootstrapped distribution of estimates for the site level random effect from M4 substantially overlapped 0, while that for M3 was well above zero (Fig. S1). Further supporting our interpretation of this term, a KS-test of the difference between these bootstrapped distributions was highly significant (*p* < 10^−10^).

## Discussion

Our results demonstrate that bees vary in their floral fidelity and that tongue length explains a large part of this variation. Bees with shorter tongues move between plant species (floral infidelity) more often than bees with longer tongues. We did not find significant variation in the response of bee species to a reduction in interspecific competition, but rather saw a guild-wide reduction in floral fidelity in response to the removal of the dominant bee species (following Brosi and Briggs 2013). Finally, our results suggest that tongue length of the most abundant bee species, a site-level attribute, explains much of the site-to-site variation in pollinator foraging behavior. In particular, we found that as the tongue length of the most abundant bee (i.e. the species that was experimentally removed) increases, the site level foraging fidelity decreases (Fig. 2).

We found that bumble bee species vary in the degree to which they move between different plant species within a single foraging bout, and tongue length explains much this variation. Some suggest that long tongued bees should exhibit broader resource usage patterns because their traits permit them access to a wider range of flower types (Santamaría and Rodríguez-Gironés 2007). In contrast, short-tongued bees should act as specialists, with a more restricted range of resource use options, rarely able to access the nectar at the base of the flowers with long corollas (Harder 1985, Graham and Jones 1996). Our results suggest the opposite pattern, that shorter tongue bees are more labile with their foraging patterns and on average move between plant species within a single foraging bout more often than longer tongue bees. We suggest the following interpretation: because long tongues enable bees to access flowers with better rewards (e.g. long-corolla flowers) and maintain a monopoly on those rewards, they may have less incentive than short-tongued species to move between plant species while foraging. While longer tongue bumble bees are capable of foraging on flowers with short corollas (Heinrich 1976, Plowright and Plowright 1997) it would provide less energetic gain, making behavioral plasticity less profitable (Inouye 1980). In contrast, the shorter-tongued bees in our system tend to have smaller bodies and are more likely to depend on resources within a more restricted foraging range (Westphal et al. 2006). These limitations could favor a labile foraging habit, with shorter tongue bees constantly assessing the resource availability and competitive context in their community. As such, shorter-tongued bees more readily switch between plant species.

We found an overall reduction in floral fidelity across sites after the removal of the most abundant bee species. Our results build on the findings of Brosi and Briggs (2013), reaffirming a guild-wide reduction in floral fidelity in response to a reduction in interspecific competition. This study adds an additional two years, 8 sites and 165 bee individuals to our previous study, confirming that the guild-wide results found in Brosi and Briggs (2013) are robust.

Variation in bumble bee floral fidelity is largely explained by the tongue length of the most abundant (i.e. removed) bee species in each site. This means that pollinator foraging behavior is context dependent and is determined (at least in part) by the most abundant bee species. In general, short tongue bees exhibit lower floral fidelity than long tongued bees but when they are in a site that has a long tongue bee removal, their reduction in floral fidelity is magnified.

Bumble bees are large bodied insects that require many floral resources to keep their colony growing throughout the (often short) growing season. As such, we might expect strong competition between these species, and tongue length, arguably one of the traits most relevant for resource acquisition, could dictate how resources are partitioned within a community, ultimately driving the assembly of bumble bees within communities (Heinrich 1976, Harmon-Threatt and Ackerly 2013). Pyke (1982) proposed that bumble-bee species with similar tongue lengths could not exist in his altitudinal alpine transects presumably because the bees compete for floral resources. But later studies did not support this pattern (Ranta 1982, Goulson et al. 2008), and, as in our study, found that bumble bee species with similar tongue lengths co-ocurred within a community. This has left researchers to wonder if the coexistence of many bee species with substantial overlap in their life history requirements is possible because bumble bee species compete for something other than flower resources (i.e. nesting sites) allowing so many similar species to co-occur (Goulson et al. 2008). Our study suggests that in our system, bumble bees do in fact compete for floral resources and that longer tongue bees seem to elicit competition that is experienced across the range of trait values seen in our sites (see Table 1). The willingness of short tongue bees to exhibit behavioral plasticity may allow for such a large number of seemingly similar bee species to coexist in a community (Valdovinos et al. 2016). Future work should examine the extent to which this plasticity is adaptive and assess the fitness costs (or benefits) that may result from the willingness to switch floral resources in response to a reduction in interspecific competition.

Most pollinators are generalist foragers that can switch between plant species within a single foraging bout (Waser et al. 1996, Brosi and Briggs 2013). When pollinators move between plant species, they can transfer heterospecific pollen to plant stigmas which in turn can reduce plant reproduction (Morales and Traveset 2008, Mitchell et al. 2009, Ashman and Arceo-Gomez 2013, Briggs et al. 2016). We found an effect in which competition from a long tongued bee changes the foraging behavior of the rest of the bees in a way that could be detrimental to plant reproduction. From a plant's perspective, not only do short tongue bees exhibit behavior that likely results in the transfer of heterospecific pollen, but when short tongue bees are in communities in which a longer tongue bee is most abundant, they exhibit even greater floral infidelity, making the likelihood of heterospecific pollen deposition even greater (see Fig. 2).

Pollinator species are on the decline globally (Potts et al. 2010). Bumble bees in particular are experiencing population range contractions due to climate change (Kerr et al. 2015) as well reductions in abundance due to disease (Cameron et al. 2011), agricultural intensification and pesticide use (Goulson et al. 2015). In a changing world where we are likely to experience an emergence of new interactions, exploring how the competitive landscape shapes foraging plasticity will help us generalize to other plant pollinator systems and begin to better predict the functional implications of competitive interactions.

## Acknowledgments

We thank P. Caradonna and A. Iler for constructive comments on the manuscript. P. Humphrey provided invaluable assistance with data analysis. Thanks to G.Gilbert and I. Parker for numerous conversations about ideas and input on drafts. Thanks also to the the Rocky Mountain Biological Laboratory staff, especially J. Reithel and I. Billick, provided key research and logistical support. P. Brenes-Coto, M. Hensley, E. Kerchner, A. Cooke and K. Niezgoda, L. Anderson, J. Brokaw, T. Lamperty, F. Oviedo, R. Perenyi, L. Thomas, L. Atalla, A. Delva, D. Picklum and K. Webster provided field assistance. This work was funded by US National Science Foundation Grants DEB-1120572 (to B.J.B.), DEB-1406262 (to H.M.B) and DBI- 1034780, OIA-0963529, and DBI-0753774 (to I. Billick), the Rocky Mountain Biological Laboratory (B.J.B. and H.M.B.), Emory University (B.J.B.), and University of California, Santa Cruz (H.M.B.).

## Supplemental Information

**Figure S1:**
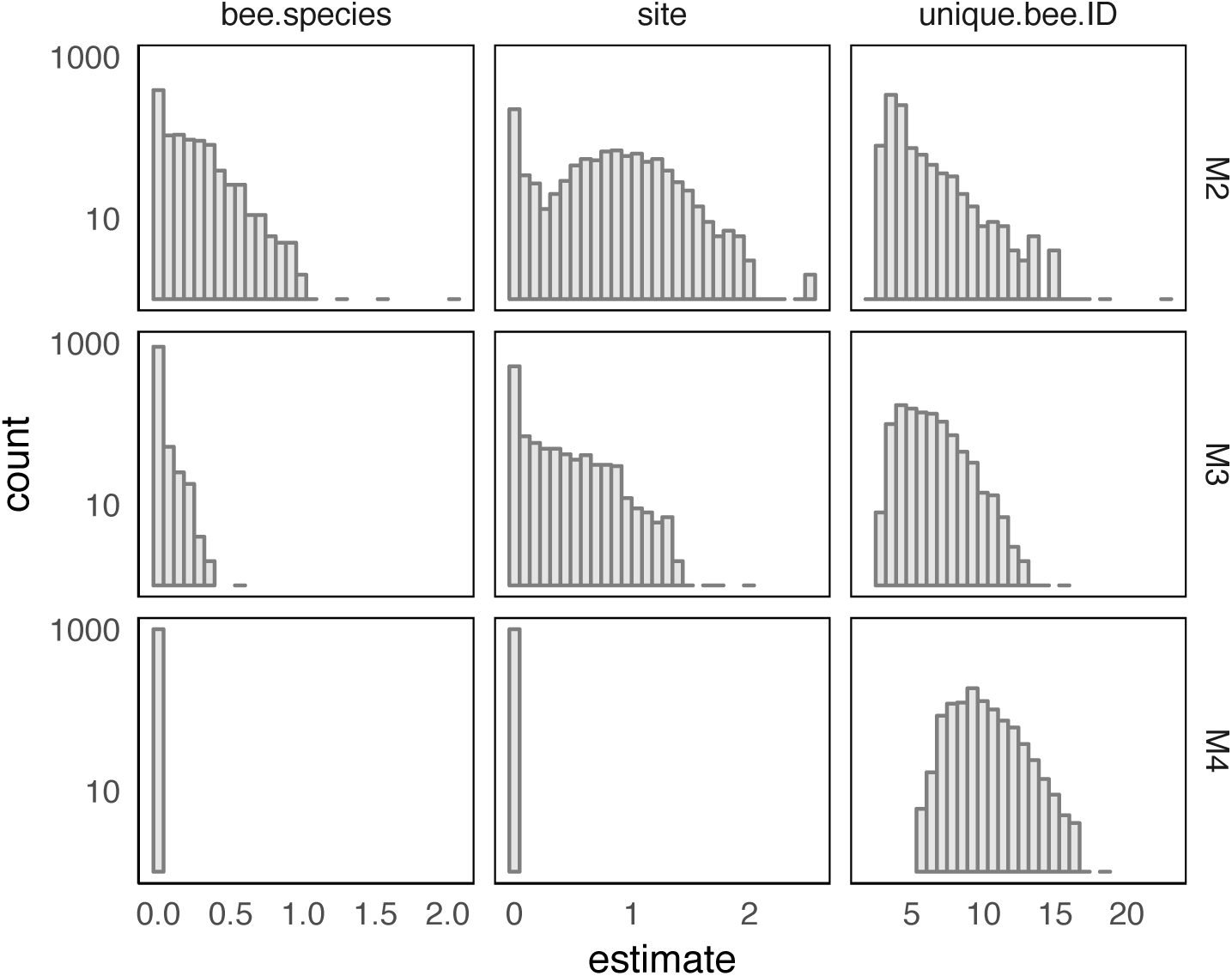
Parametric bootstrap estimates for variance terms. To evaluate the significance of the reductions in estimated random effects variances, we conducted parametric bootstrapping of models M2, M3, and M4 (1000 replicates each) and compared the distribution of estimates for σ^2^(M2) and σ^2^(M3) for the bee species random intercept (‘bee.species’), as well as σ^2^(M3) and σ^2^(M4) for the site-level random intercept (‘site’). Plotted are the distribution of estimates for each variance term in the models, with the individual level random effect (‘unique.bee.ID’) added for comparison.

**Figure S2:**
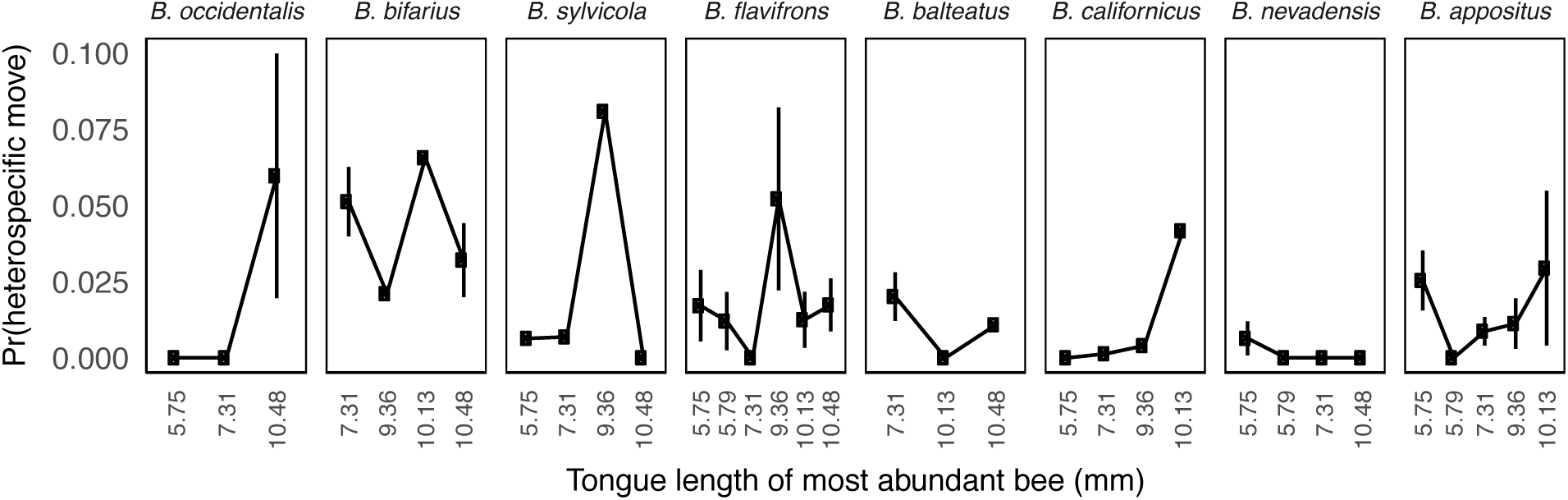
Trait based response to pollinator removals. Heterospecific move probability for each bee species as a function of trait-based competitive context (i.e. tongue length of the most abundant bee species at that site). Plotted are data from the Control condition only to reveal standing variation in foraging behavior in relation to site-level variation in the abundant bee species. Plotted are means and 95% CIs calculated on the basis of pooled counts for each bee species for a given context, as in Figure 1.

